# Acetylcholine-dependent phasic dopamine activity signals exploratory locomotion and choices

**DOI:** 10.1101/242438

**Authors:** J. Naudé, S. Didienne, S. Takillah, C. Prévost-Solié, U. Maskos, P. Faurej

## Abstract

Dopamine neurons from the Ventral Tegmental Area (VTA) switch from tonic to phasic burst firing in response to reward-predictive cues and actions. Bursting is influenced by nicotinic acetylcholine receptors (nAChRs), which are not implicated in reinforcement learning, but rather in exploration and uncertainty-seeking. The leading model assigns these functions to tonic dopamine firing. To investigate this paradox, we recorded the activity of VTA dopamine neurons during a spatial decision-making task. When reward was certain, mice adopted a stereotyped behavior, and dopamine neurons signaled reward. When confronted with uncertain rewards or a novel environment, mice exhibited exploration. Modulation of phasic, but not tonic, dopamine activity predicted uncertainty-seeking and locomotor exploration. Deletion of nAChRs disrupted the influence of uncertainty and novelty on dopamine firing and behavior, sparing reward signaling and learning. Hence, nAChR modulation of dopamine neurons can influence cognitive functions on a short timescale, through the modulation of phasic, synchronous bursting.

## INTRODUCTION

VTA dopamine neurons play a major role in motivating goal-directed behaviors and reinforcing behaviors leading to reward (Glimcher, 2011; Schultz, 2007; Sutton and Barto, 1998). These cells encode a “reward prediction error”, the difference between predicted and actual reward, which can be used to update the expected (mean) value of stimuli and actions (Glimcher, 2011; Schultz, 2007). Dopamine neurons signal a reward prediction error through a switch from tonic (regular single-spike firing) to phasic (synchronous bursting) firing (Faure et al., 2014; Grace et al., 2007; Schultz, 2007). The bursting pattern in dopamine cells depends on the balance between glutamatergic and GABAergic inputs (Lobb et al., 2010; Paladini and Roeper, 2014; Zweifel et al., 2009), but also critically on cholinergic modulation (Dani and Bertrand, 2007; Faure et al., 2014; Grace et al., 2007; Mameli-Engvall et al., 2006). Understanding how mesopontine acetylcholine impacts decision-making through the modulation of firing in dopamine cells is of utmost importance, as dysregulations of these neuromodulatory systems are implicated in major psychological diseases such as schizophrenia, tobacco addiction or Parkinson’s disease (Dani and Bertrand, 2007).

In particular, nicotinic acetylcholine receptors (nAChR) containing the β2-subunit constitute major determinants of dopamine neurons firing pattern. In anesthetized mice lacking the nAChR β2-subunit, dopamine cells lack spontaneous bursting (Mameli-Engvall et al., 2006; Maskos et al., 2005; Naudé et al., 2016). Contrasting with the role of dopamine cell bursting in reinforcement learning (Faure et al., 2014; Grace et al., 2007; Schultz, 2007), blocking or deleting β2-containing nAChRs (β2*nAChR) does not affect reward-guided learning (Naudé et al., 2016), motivation to work for reward (Yeomans and Baptista, 1997), or choices between differently sized rewards (Serreau et al., 2011). Nonetheless, VTA β2*nAChRs are implicated in complex cognitive processes such as exploration (Granon et al., 2003; Maubourguet et al., 2008; Naudé et al., 2016) and uncertainty processing (Naudé et al., 2016), which are believed to depend on tonic fluctuations (Beeler et al., 2010; Fiorillo et al., 2003; Frank et al., 2009; Humphries et al., 2012; Niv, 2007; Schultz, 2007).

There is thus an apparent discrepancy between the proposed behavioral roles of VTA nAChRs (exploration, but not reward), the kind of dopamine firing that they are thought to control (switch from tonic to phasic) and the respective roles assigned to phasic (reward) and tonic (exploration) dopamine. In other words, direct evidence of how nAChRs affect dopamine firing and function during exploration or uncertainty-processing tasks is still lacking. Here we assessed dopamine encoding of uncertainty, novelty and exploration in a self-paced task, with no stimulus signaling the beginning of a trial, in order to give the mice full control over whether to exploit reward or explore the open-field. We found that dopamine neurons encoded expected reward and motivation for both reward-directed decisions and locomotion, i.e., exploitation, in purely predictable contexts. Importantly, we also found that the phasic activity of dopamine neurons summed uncertainty with reward, to signal motivation to explore in uncertain or novel environments. We then disrupted β2*nAChRs to relate alterations in uncertainty processing and exploration with dopaminergic signaling, and show that dopamine neurons in β2^−/−^ mice did not encode uncertainty nor exploration bonuses, concomitantly with the disappearance of uncertainty-seeking and exploratory locomotion.

## RESULTS

### β2*nAChRs control spontaneous, synchronous bursting in dopamine neurons

We recorded the spontaneous firing from putative dopamine neurons (pDAn) in the VTA from WT (n=87 from 16 mice) and mice deleted from the β2-subunit of the nAChRs (β2^−/−^, n=80 from 12 mice) awake mice at rest, using extracellular poly-electrodes (Fig. 1a). All neurons met the criteria used to identify dopamine cells in vivo (Roesch et al., 2007; Takahashi et al., 2016) (Fig. 1b, Supplementary Fig. 1, Methods). Spike trains were decomposed based on inter-spike intervals (ISI) into phasic bursting (corresponding to shorter ISI, see Methods) and tonic firing patterns (Fig 1c). On average, pDAn from β2^−/−^ mice fired at a lower frequency (WT=4.96Hz, β2^−/−^=4.11Hz, U^(87,80)^=8171, p=0.006, Fig. 1d) than that from WT mice. However, there was no difference in the proportion of spikes within bursts (%SWB) between the two genotypes (WT=40.5%, β2^−/−^=36.5%, U_(87,80)_=7705, p=0.2). This contrasts with previous studies using anesthetized β2^−/−^ mice, in which DA neurons lack bursting (Mameli-Engvall et al., 2006; Naudé et al., 2016), suggesting a different control of DA firing in awake mice. Synchronous bursting activity among dopamine neurons causes a substantial larger dopamine release in target structure (Grace et al., 2007) than asynchronous activity. We thus next measured basal synchrony between raw spike trains (all spikes), and also assessed synchrony between bursting events (all spikes within bursts), bursting onsets (first spike of bursts) and tonic spiking (spikes outside bursts). We found that bursting onsets were spontaneously synchronized (Kreuz et al., 2013)- in the sense that synchrony was higher than chance level in WT (Eshel et al., 2016; Joshua et al., 2009), but not in β2^−/−^ animals (p<0.006, KS-test, Fig. 1e, Supplementary Fig. 2). Hence, in awake mice, bursting events are spontaneously synchronized among VTA DA neurons, and their synchronization is under the control of β2*nAChRs.

**Figure 1.**
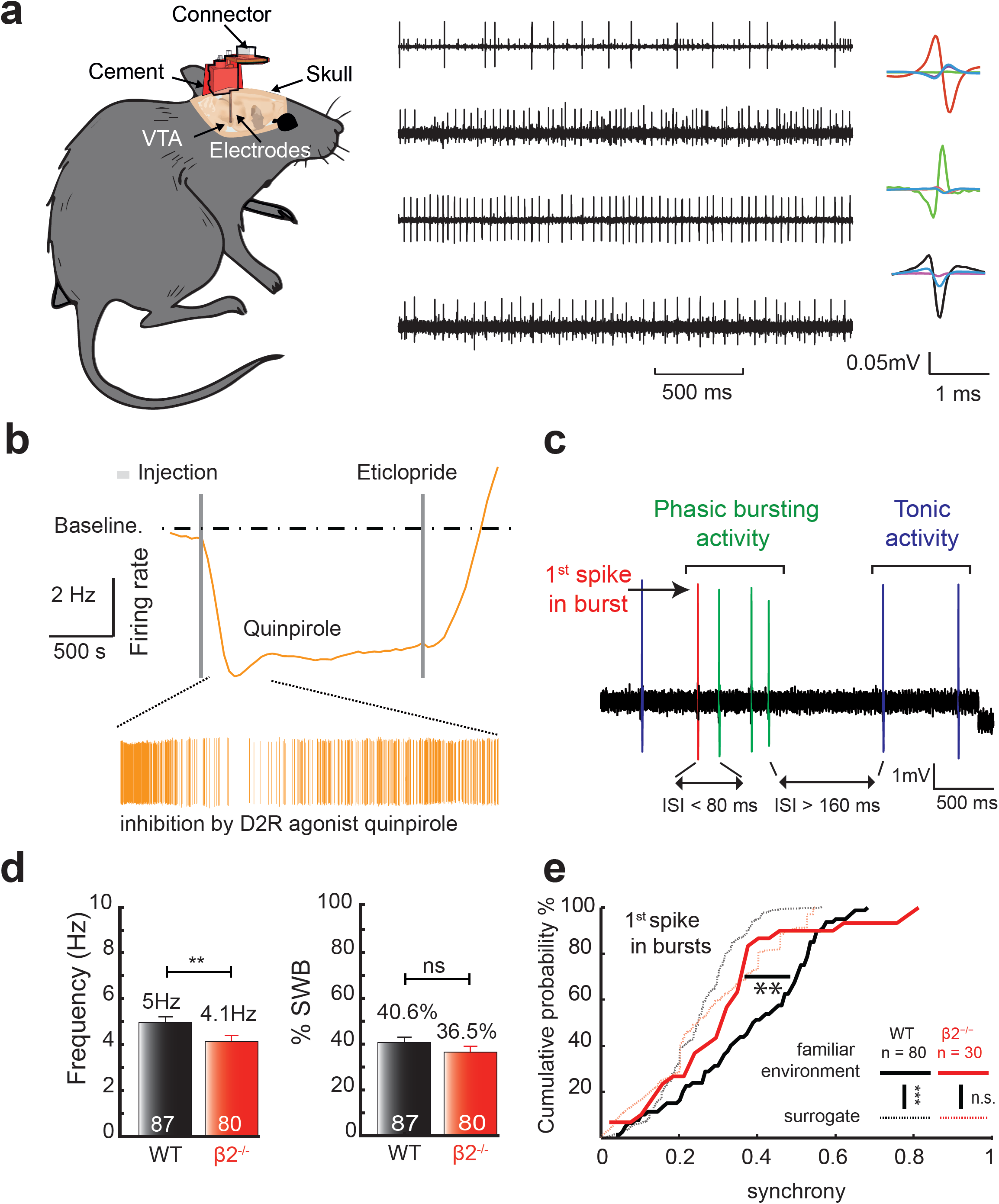
β2*nAChRs control synchronous bursting in dopamine neurons. (**a**) Schematic of in vivo implantation (left). Filtered extracellular recordings in the VTA (middle) and average spike waveforms from simultaneously recorded neurons (right). (**b**) Pharmacological confirmation of the dopaminergic nature of recorded neurons. Neurons were considered dopaminergic if inhibited by intra-peritoneal injection of D2R agonist quinpirole followed by reactivation by D2R antagonist eticlopride. (**c**) Decomposition of spike trains into phasic bursting and tonic firing patterns. Bursts start with an interspike interval (ISI) less than 80ms and stop when the ISI become greater than 160ms. All spikes within bursts were labeled “phasic” while spikes outside bursts were labeled “tonic”. (**d**) Firing frequency (left) and %SWB (right) of pDAn from WT (n=87 neurons from 16 mice) and β2^−/−^ mice (n=80 neurons from 12 mice). (**e**) Cumulative density of pairwise synchrony between burst onsets in WT (black) and β2^−/−^ (red) mice. Solid lines correspond to spontaneous synchrony in familiar environment. Dashed lines correspond to surrogate data (shuffling of neurons pairs). Burst onsets from WT mice show increased synchrony over surrogate (for WT mice, p<0.001; for β2^−/−^ mice, burst onsets: 0.14; KS tests) and synchrony between burst onsets was different in WT and β2^−/−^ mice (p=0.006, KS test).

### Synchronous bursting emerges with the learning of a self-paced task

Dopamine cells signal reward by a phasic activation, i.e. a time-locked increase in synchronous bursting activity. We thus assessed how the involvement of β2*nAChRs in synchronous bursting can affect phasic dopamine activity related to reward processing. Phasic activity from dopamine cells is usually conceptualized as a reward prediction error (Glimcher, 2011; Sutton and Barto, 1998): δ(*t*)= *R*(*t*)+γ*V*(*s_t_*_+1_) − *V*(*s_t_*), where δ(*t*), the reward prediction error, is assumed to correspond to phasic dopamine activity, *R*(*t*) to the actual reward eventually obtained, γ*V*(*s_t_*_+1_) to the prediction of the value of the next choice (temporally-discounted by a factor γ) and *V*(*s_t_*) to the expected value of the current state of the animal. We determined whether the activity from VTA pDAn (126 from 12 WT and 103 from 10 β2^−/−^ mice) is compatible with a reward prediction error, while mice performed a spatial version of the multiarmed bandit task (Naudé et al., 2016) (Methods). In an open-field, mice learned to associate three explicit locations with intracranial self-stimulation (ICSS (Carlezon and Chartoff, 2007)) (Fig. 2A). Because two consecutive rewards cannot be obtained at the same location, mice learnt to alternate between them. When faced with certain rewards, both WT and β2^−/−^ mice learned the task (F_(9,13)_=118.18, p<0.001) and performed similarly (F_(1,22)_=2.2, p=0.89, Fig. 2a).

**Figure 2.**
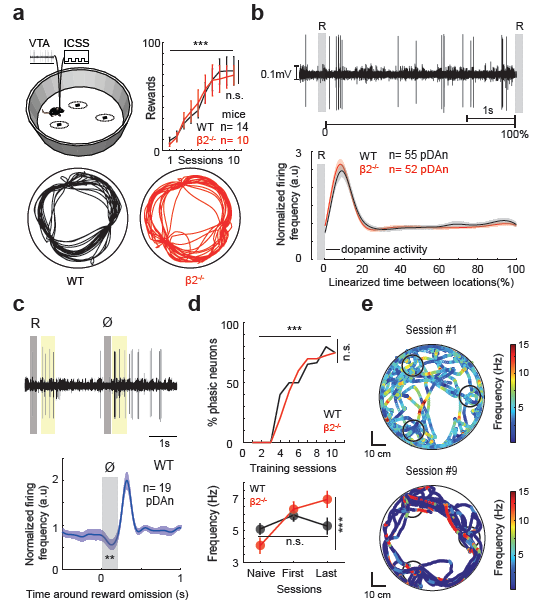
Reinforcement learning reorganizes dopaminergic firing towards the self-initiation of actions with learning, independently from β2*nAChRs. (**a**) Top: multi-armed bandit task using ICSS (left) and number of ICSS rewards against learning sessions (right). Bottom: representative trajectories for WT and β2^−/−^ mice after learning. (**b**) Top: extracellular recordings between two ICSS (grey boxes) after learning. Bottom: normalized firing frequency against normalized time between two ICSS for WT (n=55 neurons from 13 mice) and β2^−/−^ mice (n=52 neurons from 10 mice). (**c**) extracellular voltage (top) and normalized firing frequency (bottom) during the omission of an ICSS, showing examples of early bursting related to next reward (yellow boxes). (**d**) Top: percentage of pDAn displaying movement onset-related activity against learning sessions. Bottom: average firing frequency throughout learning (naive: open-field without ICSS, first: sessions 1–5, last: sessions 6–10). (**e**) Representative examples of animal trajectory during a session, with corresponding instantaneous firing frequency (color coded), in the first session (top) and late session (bottom), illustrating correlated reorganization of behavioral and spiking dynamics through learning.

After learning, pDAn emitted bursts of action potentials early in the trials (Fig. 2b), henceforth called “early bursting”. Early bursting emerged in both WT and β2^−/−^ animals, suggesting that the phasic increase in dopamine activity at actions onset (Jin and Costa, 2010; Wassum et al., 2012) is β2*nAChRs-independent. Early bursting was characterized by a series of features, in both WT and in β2^−/−^ mice, compatible with the encoding a reward prediction error occurring during the self-initiation of an action that constitutes the first predictor of the next reward (Sutton and Barto, 1998; Wassum et al., 2012). First, early bursting was not triggered by the electrical stimulation, since it occurred when ICSS reward was unexpectedly omitted (Fig. 2c), in the first trials of the probabilistic setting used later on (see below). Hence, early bursting appeared to predict the next reward, rather than signaling the previous one. Second, during reward omissions (Fig. 2c), pDAn activity decreased at the time of the expected reward (t_(18)_=-3.07, p=0.007). This result is also consistent with dopamine cell signaling a reward prediction error (Glimcher, 2011; Schultz, 2007; Sutton and Barto, 1998), as the actual reward is lower than expected: in the first omission trials, animals expect a reward but do not obtain it (*R*(*t*)=0 so that *R*(*t*) − *V*(*s*_*t*_)<0). Finally, the percentage of pDAn displaying a bursting activity significantly larger than baseline firing at any time between two ICSS increased over learning sessions in WT mice (χ2=7.81, p=0.01, Fig. 2d). The increase in phasic activity was also apparent in pDAn from β2^−/−^ animals (proportion of pDAn displaying phasic activity not different from WT: χ2=0.05, p=0.82). This gradual increase in the total number of VTA dopamine cells displaying phasic bursting activity is consistent with the emergence of a positive reward prediction error during learning.

### Reorganization of firing with learning

We next analyzed how this change in dopamine firing at the single-trial scale impacted the average activity of dopamine neurons. Completion of learning induces the emission of bursts by dopamine cells at each trial, but also corresponds to an increase in the number of trials, the combination of these two features thus suggests an overall increase in pDAn firing frequency. However, this was not observed: even though the %SWB and burst synchrony increased throughout learning (%SWB: F_(2,118)_=49.78, p<0.001, synchrony: F_(2,141)_=63.1, p<0.001, Supplementary Fig. 3), the average firing frequency remained constant (F_(2,118)_=1.3, p=0.28, Fig. 2d below). Hence early bursting in dopamine neurons (Jin and Costa, 2010; Wassum et al., 2012), reflecting reward prediction, does not rely on additional spikes, but rather on a dynamical reorganization towards synchronous bursting activity (Fig. 2e). In β2^−/−^ mice, the increase in phasic activity in pDAn was however accompanied by an increase in average firing frequency (genotype x learning interaction F_(2,216)_=4.81, p=0.01; learning effect in β2^−/−^: F_(2,98)_=9.15, p<0.001), and pairwise synchrony between pDAn bursting increased less in β2^−/−^ than in WT mice (genotype: F_(1,232)_=11.85, p < 0.001; genotype x learning interaction: F_(1,2)_=3.47, p = 0.03, Supplementary Fig. 3). This indicates β2*nAChRs are not only implicated in the spontaneous synchrony among DA neurons (Fig. 1), but also in the re-organization of collective dynamics of the VTA upon learning.

### Dopamine neurons signal exploitation of rewards in predictable environment

Reinforcement models also state that after learning, dopaminergic reward prediction does not play any further role in ongoing behavior (Glimcher, 2011; Sutton and Barto, 1998). Recent studies have shown that phasic dopamine can also encode for kinetic variables, but this mainly concerns dopamine neurons from the Substantia Nigra pars compacta (SNc)(Barter et al., 2015; Howe and Dombeck, 2016). We thus asked whether pDAn from the VTA encode for motor variables. In our setup, early bursting occurred during the animal’s dwelling time, before movement onset towards the next location (Fig. 3a). When assessing the correlations between early bursting and kinetic variables, we did not find any relation with immediate behavior (e.g. instantaneous speed or acceleration just following early bursting, Supplementary Fig. 4), as found in studies interested in the role of the SNc in motor control (Barter et al., 2015; Howe and Dombeck, 2016). However, we observed that a higher frequency during early bursting corresponded to a shorter trial duration, i.e. the time needed by the animal to reach the next rewarded location (Fig. 3b). Sorting the trials by increasing “early bursting” frequency (i.e. frequency during the first second of the trial, see Methods) revealed that it correlated with the time-to-peak speed (Fig. 3c) and, more precisely, with the ratio of maximum speed over the time-to-maximum speed (Fig. 3d, WT: 62.5% modulated cells, median R^2^: 0.54; β2^−/−^: 76.0% modulated cells, median R^2^: 0.58). This ratio relates to the directness of the animal towards the goal: a high peak speed, a strong acceleration, a direct path or a short dwell-time all increase this measure. Since ICSS reward was equivalent at all locations (and thus, the reward prediction would be the same for all trials), our findings suggest that dopamine phasic activity encodes a locomotor signal related to the whole action sequence (from start to goal)(Wassum et al., 2012), on top of a reward prediction error. Moreover, the proportion of cells showing directness-related firing was similar in WT and β2^−/−^ mice (p=0.21, KS-test), indicating that VTA dopamine signaling of locomotor activity is β2*nAChRs-independent in the context of exploitation.

**Figure 3.**
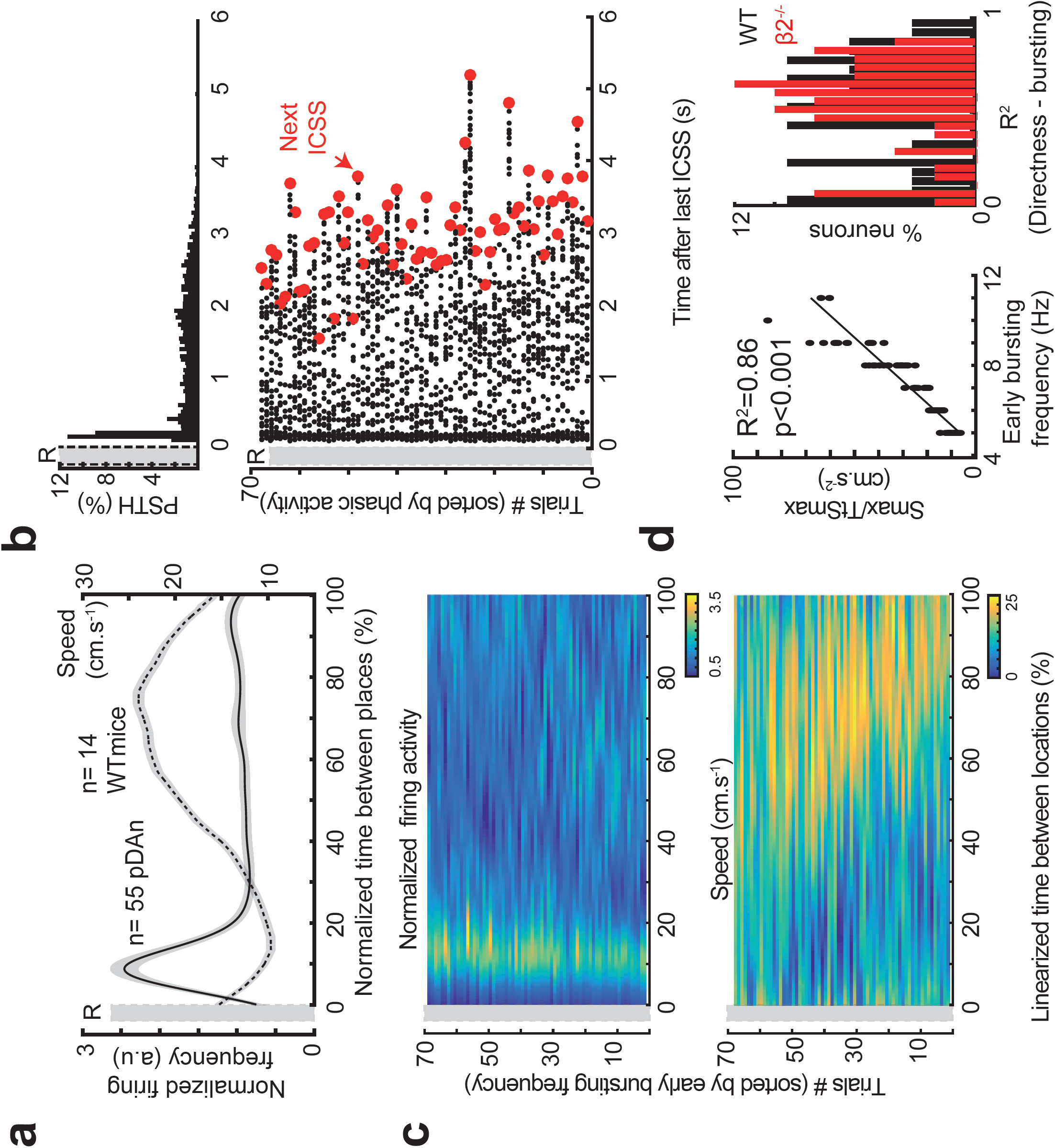
Dopamine signaling of reward exploitation predicts directness towards the goal, independently from β2*nAChRs. (**a**) Normalized firing frequency (solid lines) and instantaneous speed (dashed lines) against normalized time between two ICSS for WT mice. (**b**) Bottom, raster plot of action potentials, sorted by increasing phasic activity of pDAn. Each line is a trial between two ICSS locations, each black dot is an action potential, and red dots represent the end of the trial. Top, histogram of the raster plot. (**c**) Normalized firing activity (top) and speed profiles (bottom) corresponding to the raster plot in (A), against linearized time between two ICSS locations, sorted by increasing phasic activity of pDAn. (**d**) Representative example of the correlation between firing frequency at movement onset and the peak speed over time-to-peak speed ratio (left) and distribution of the coefficient of determination (R^2^) for the correlation in (**c**) for all neurons (right).

### Encoding of uncertainty by phasic bursting in dopamine neurons

We next investigated pDAn signaling in decision-making under uncertainty, using a probabilistic setup, in which the three rewarded locations are respectively associated with different probabilities to receive ICSS (Fig. 4A, p=100%, p=50% and p=25%). The three binary choices (here, 25% vs. 50%, 25% vs. 100% and 50% vs. 100% reward probabilities) allow characterizing the influence of two co-varying parameters (reward mean, which corresponds to *p* and uncertainty, defined as variance *p(1-p)*) on decisions (Naudé et al., 2016). Mice visited overall the locations associated with higher ICSS probability more often (Fig 4b left), indicating a choice behavior. However, when starting from the 25%-reward probability location, they chose the 50% reward as much as the 100% reward (U=35, p =0.49, Wilcoxon test, Fig 4b middle), suggesting that mice are irrationally attracted by the 50% reward. We investigated this preference by fitting the binary choices with a computational model that takes into account the relative influences of reward mean and reward uncertainty on decisions (see *Methods*). This model-based analysis indicates that WT mice chose as if they assigned a positive value to the reward uncertainty (a “bonus” value added to expected reward) (Kakade and Dayan, 2002; Naudé et al., 2016) of the goal (Fig 4B right), which is maximal for 50% reward probability (Fig. 4a). In other words, mice choices appear suboptimal because they explore actions with uncertain consequences.

**Figure 4.**
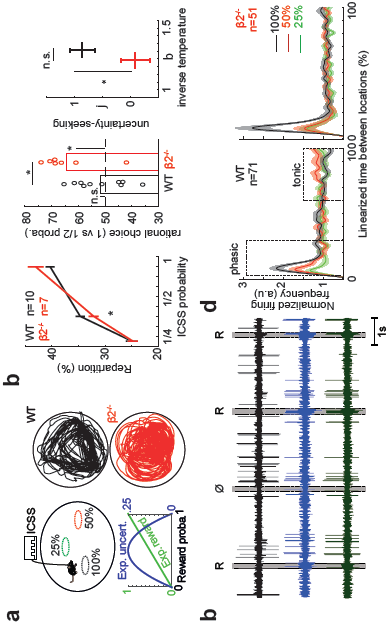
β2*nAChRs bias choice of uncertain rewards and its encoding by dopamine neurons. (**a**) Left schematics: in probabilistic sessions, each location was associated with a distinct ICSS probability. Right: Representative trajectories for WT and β2^−/−^ mice. (**b**) Left: proportion of choices against reward probability. Middle: proportion of irrational choices of uncertain outcomes, defined as choosing the location associated with 50% reward probability against 100% reward probability. Right: uncertainty bonus versus inverse temperature, fitted from the choices. (**c**): Extracellular recordings when the animal received (R) or not (ø) an ICSS. Dashed red boxes indicate phasic activity (P). (**d**): normalized firing frequency against normalized time between two locations for WT (left, n=71 neurons from 13 mice) and β2^−/−^ mice (right, n=51 neurons from 10 mice), sorted by reward probability at the goal. Dashed boxes delineate activity modulated by reward probability (early bursting and late ramping activity) and affected by the genetic deletion of β2*nAChRs.

We next recorded pDAn firing during this task (Fig 4c) to assessed whether the bonus value of uncertainty was encoded by pDAn neuron. pDAn firing activity was sorted according to the reward probability of the goal during direct trials (duration shorter than 5s, average median duration of trials =4.84±0.39s, mean±s.e.m. in WT mice) in WT and β2^−/−^ mice. Averaging trials with different durations by linearizing the time between two consecutive locations (Fig. 4D, Supplementary Fig. 5 displays all neurons) revealed that (1) early bursting, but also late, “ramping” activity, seemed modulated by reward probabilities (boxes in Fig. 4D) and (2) these dynamics seemed altered by the genetic deletion of β2*nAChRs (Fig. 4D left versus right). We thus centered the analysis of early bursting on the first 500ms of trial onset (Methods). Early bursting was sensitive to the expected reward probability at the goal (F_(2,187)_=5.53, p=0.005, Fig. 5A), consistent with a reward prediction (Glimcher, 2011; Schultz, 2007). In addition, encoding of uncertain rewards (50%) relative to certain rewards (100%) by pDAn early bursting activity predicted the bonus (i.e. the positive value) mice assigned to uncertainty (R^2^=0.47, p=0.03, Fig. 5B). This suggests that the phasic activity of VTA dopamine neurons integrates an uncertainty bonus together with expected value (Kakade and Dayan, 2002; Lak et al., 2014).

**Figure 5.**
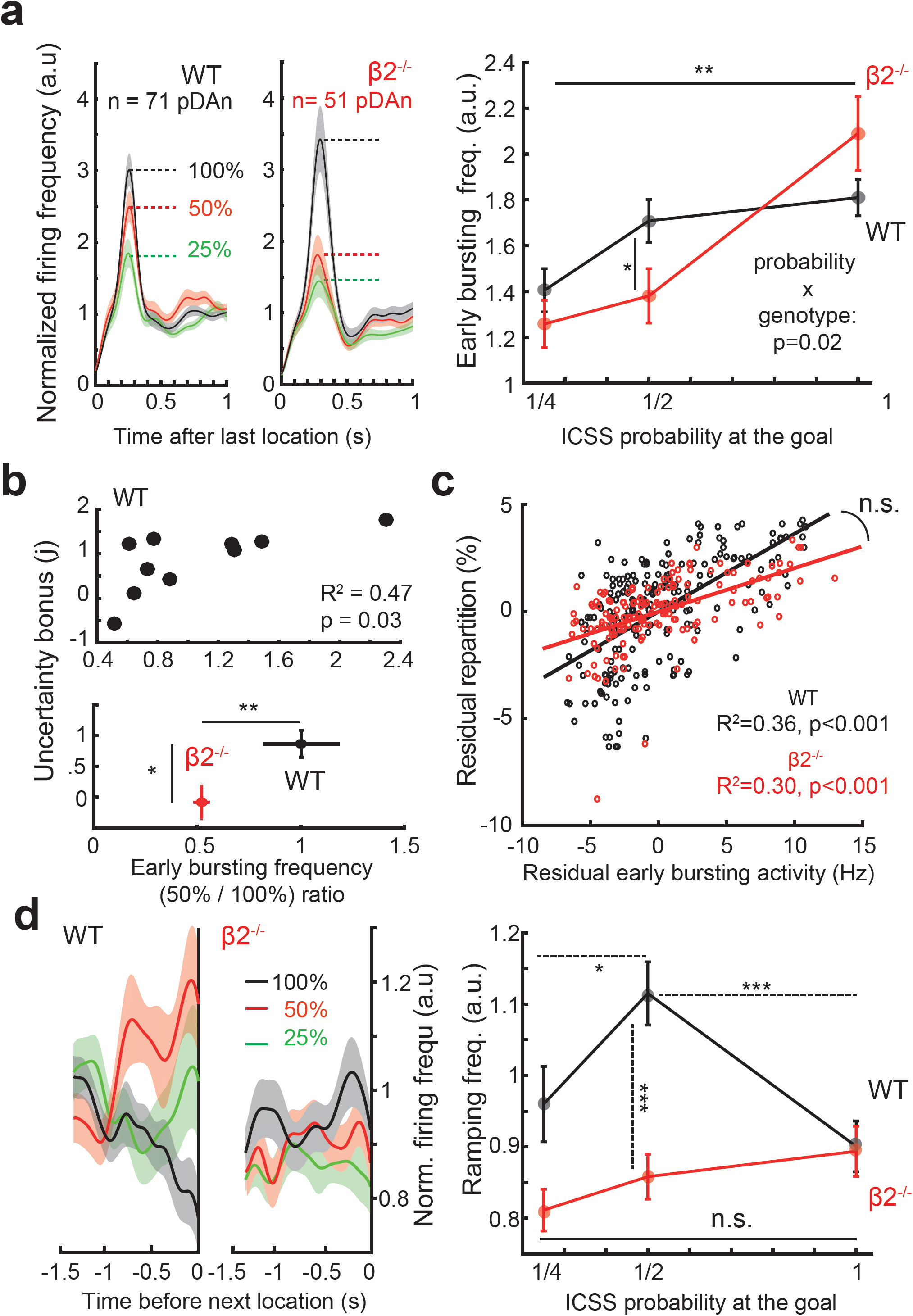
Valuation of uncertain rewards is predicted by phasic, but not tonic, activity of dopamine neurons. (**a**) Normalized firing frequency after the last location for WT (left) and β2^−/−^ (middle), and early bursting activity against reward probability at the goal (right). (**b**) Uncertainty bonus against encoding of uncertainty by pDAn early bursting (phasic activity related to 50% versus 100% reward probability). Top: each dot represents a WT mouse. Bottom: group averages for WT and β2^−/−^ mice. (**c**) Residual choices against residual early bursting activity. (**d**) Normalized firing frequency before the next location for WT (left) and β2^−/−^ (middle), and firing frequency during ramping activity against reward probability at the goal (right).

Moreover, we found a significant correlation between the residuals of pDAn activity and of the next choices of the animal (R^2^=0.36, p<0.001, Fig. 5C), after subtracting the effect of reward probability on both variables (Varazzani et al., 2015) (Methods). For instance, in a given session, if a mouse chose a rewarded location more than the group average, pDAn from this mouse displayed an activity corresponding to this location’s reward probability higher than the population average. In other words, dopamine activity encodes the subjective value of the next choice rather than the objective reward attributes (e.g. reward probability). Besides, early bursting activity correlated with the peak speed/time-to-peak speed ratio, as in the certain setting, but to a lesser extent (Supplementary Fig. 6), as we discuss below when analyzing exploratory locomotion. Hence, phasic activity of pDAn signals both ongoing choices and motor control (Howe and Dombeck, 2016; Jin and Costa, 2010; Salamone and Correa, 2012). Overall, our results suggest that phasic activity in VTA dopamine cells (early bursting) incorporates different sources of value (expected reward, uncertainty-bonus, among others)(Fiorillo, 2011; Lak et al., 2014; Stauffer et al., 2014) to signal ongoing behavior.

### Uncertainty-seeking and encoding of uncertainty by dopamine neurons both depends on nicotinic acetylcholine receptors

Given the implication of β2*nAChRs in uncertainty-seeking and exploration (Maskos et al., 2005; Naudé et al., 2016), we next assessed their roles in the probabilistic task. Overall, β2^−/−^ mice chose the 50% location less than their WT counterparts (F_(1,2)_=4.19, p=0.02, repartition*genotype interaction, t_(15)_=2.31, p=0.035, Fig. 4b left). This was particularly clear in the binary choice between 50% and 100% reward, in which β2^−/−^ animals chose less the 50% alternative than WT animals (t_(15)_=2.56, p=0.02, Fig. 4b middle). This alteration in decision-making can be modeled as an absence of valuation of uncertainty (uncertainty-bonus in β2^−/−^ versus WT mice: U_(10,7)_=114, p=0.02, and not different from 0: W_(7)_=12, p=0.81, Fig 4b right, 5a right). Reward probability was differently encoded in β2^−/−^ and WT pDAn (F_(1,2)_=3.95, p=0.02, probability*genotype interaction), with a reduction of activation by uncertain rewards (t_(136)_=2.18, p=0.03, Fig. 5a) that paralleled decreased uncertainty-seeking in β2^−/−^ mice (Fig. 5b, Supplementary Fig. 7). By contrast, the correlation between residuals in early bursting and choices was the same as in WT mice (Fig. 5c), indicating that the relation between pDAn activity and ongoing choices was not altered in β2^−/−^ mice. Hence, uncertainty-seeking and encoding of uncertainty by VTA pDAn activity are both affected by β2*nAChR deletion, but not the relation between DA activity and choices.

**Figure 6.**
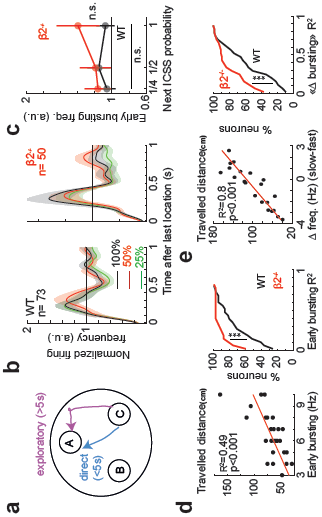
β2*nAChRs affect dopaminergic encoding of exploratory locomotion. (**a**) Schematic of the duration-based separation between direct (<5s) and long (5–10s) trials. Data presented in panels b-e were obtained during long trials only. (**b**) Normalized firing frequency over time after the last location, for WT (left) and β2^−/−^ (right). (**c**) Normalized firing frequency during early bursting activity against reward probability at the goal. (**d**) Left: example of travelled distance during the trial against firing frequency during early bursting activity. Right: Cumulative density of the correlation (coefficient of determination, R^2^) for all pDAn. (**e**) Left: example of total travelled distance against difference in bursting activity between periods of slow and fast locomotion. right: Cumulative density of the correlation (R^2^) for all pDAn.

**Figure 7.**
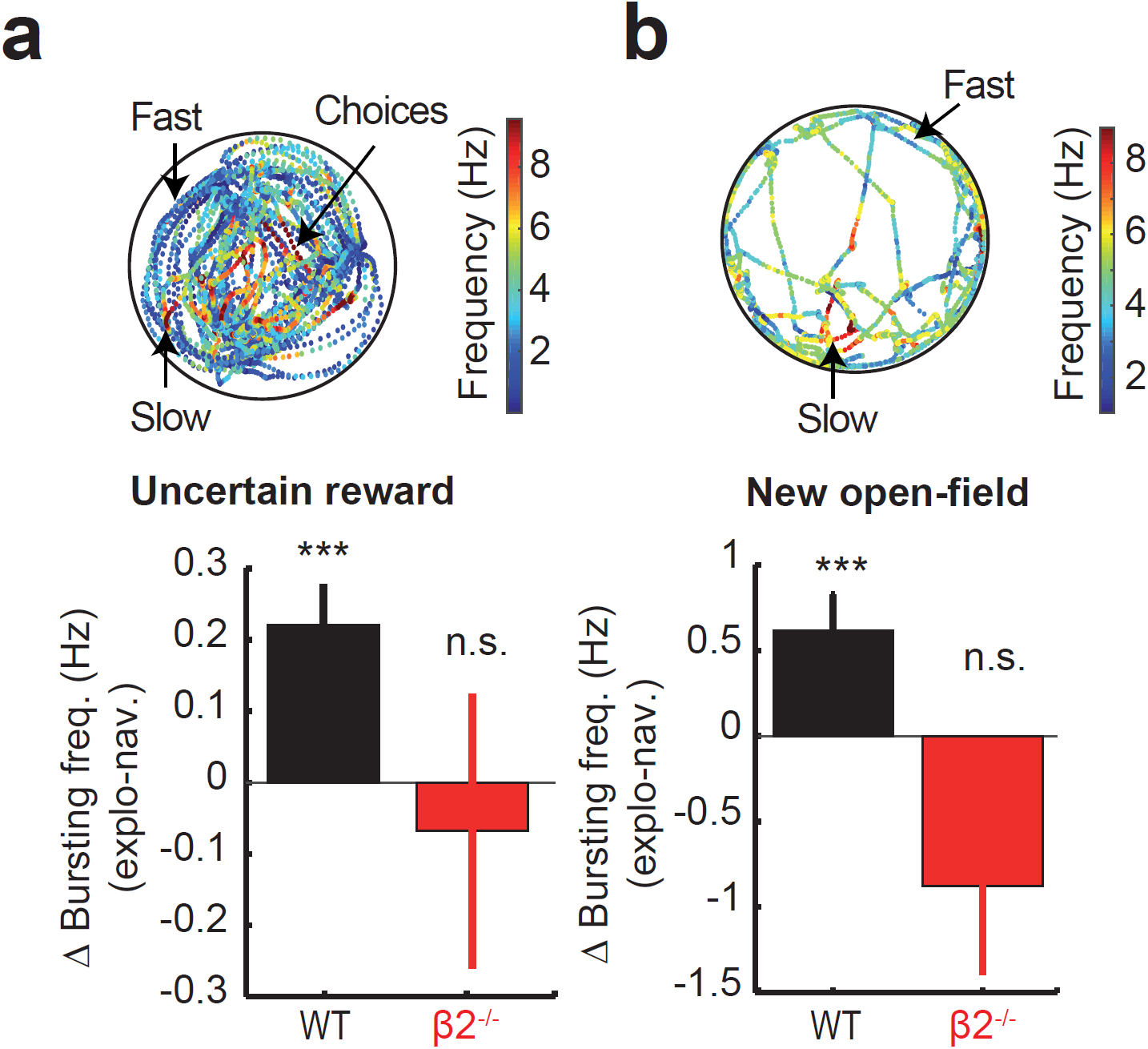
VTA dopamine neurons signal exploration induced by uncertainty and novelty. (**a, b**) Animal trajectory with corresponding instantaneous firing frequency (color coded, left) and average difference in bursting activity between periods of slow and fast locomotion (right) in a probabilistic session (**a**) and in a novel open-field without reward (**b**).

### Tonic dopamine activity encodes uncertainty but does not signal uncertainty-seeking

When no action is required from the animals, dopamine cells have been shown to encode reward uncertainty through a ramping increase in activity (Fiorillo et al., 2003), but this remains unclear in operant tasks, where uncertainty motivates the animal to explore uncertain options (Funamizu et al., 2012; Heilbronner, 2013; Naudé et al., 2016; Oudeyer, 2007). We thus sorted pDAn activity upon arrival at a location (i.e. last second before trial end, Methods) by the reward probability associated with this location (Fig. 5D left). Tonic ramping occurred before WT mice reached their goal, especially for the uncertain 50%-reward probability location (F_(2,187)_=6.36, p=0.002, 50% versus 100% encoding: t_(128)_=3.71, p<0.001). Comparison with other probability sets (Supplementary Fig. 5) indicates that pDAn ramping activity scaled with reward uncertainty. In β2^−/−^ mice, ramping was not modulated by reward probability (F_(2,137)_=1.62, p=0.20) and was different from WT mice (F_(1,2)_=4.42, p=0.01, probability*genotype interaction), indicating an implication of β2*nAChRs in uncertainty-modulation of both phasic and tonic dopamine activity (Fig. 5D right). Yet, the modulation of tonic ramping was not correlated with uncertainty-seeking behavior (Supplementary Fig. 8). This result is consistent with ramping activity appearing long after the choice, and thus not being implicated in the decision. Moreover, we observed that while early bursting activity could be considered as phasic (i.e. both bursting and synchronous), late ramping activity was mostly composed of tonic spiking (Supplementary Fig. 9). This contradicts that ramping activity might constitute a backpropagation of reward prediction error (Fiorillo et al., 2005; Niv et al., 2005). Our results thus suggest that phasic, rather than tonic, DA activity, underlies uncertainty-seeking.

### Dopamine neurons signal exploratory locomotion in uncertain or novel environments, under nicotinic regulation

Our results suggest a role of β2*nAChRs in exploration, in the framework of decisionmaking between clearly defined alternatives (here, the rewarded locations). Exploration also refers to locomotion in the rest of the open-field (Maubourguet et al., 2008) when animals wander inbetween rewarded locations in the probabilistic task, or when animals travel in a novel open-field. We thus assessed whether pDAn activity (phasic and or tonic) encodes motor variables of locomotor exploration in long trials (duration longer than 5s, Fig. 6A), during which mice display pauses and tortuous trajectories in-between rewarded locations. At the beginning of these long trials, pDAn displayed bursting activity (Fig. 6B) that did not scale with reward probability (Fig. 6C; WT F_(2,173)_=0.22, p=0.81; β2^−/−^ F_(2,117)_=0.42, p=0.66). This early bursting positively correlated with the travelled distance in WT (32.9% modulated pDAn, median R^2^=0.16) but not in β2^−/−^ mice (4.0% modulated pDAn, median R^2^=0.03, Fig. 6D). Hence, pDAn could signal distance (Fig. 6) in long trials and speed in short trials (Fig. 3). The phasic activity of DA neurons may encode the initiation and energization of a relevant motor program in both cases: speed (directness) during exploitation, and distance during exploration. This may explain why we found less directness-encoding cells in the probabilistic setting (Supplementary Fig. 6), where animals alternate between exploiting rewarded locations and exploring the rest of the open-field, than in the deterministic setup where mice only exploit rewards.

Since locomotion in long, exploratory trials in the probabilistic setting consists in a sequence of movements interspersed with pauses (Maubourguet et al., 2008), we refined the analysis of pDAn activity by distinguishing, in long trials, exploration (at low speed) from navigation (at higher speed). Periods of low speed in the open-field corresponds to rearing, scanning, sniffing and grooming (Maubourguet et al., 2008), with all these behaviors related to collecting information, except grooming that only accounts for 10% of low-speed epochs in the open-field (Maubourguet et al., 2008). During long trials, the difference between bursting activity during slow and fast epochs correlated with the total travelled distance in WT (60.3% modulated cells, example in Fig. 6E left, median R^2^=0.40, distribution in Fig. 6E right). This correlation with travelled distance was abolished in β2^−/−^ mice (10.0% modulated cells, median R^2^=0.04, Fig. 6E). Moreover, bursting activity was higher during epochs of slow exploration compared to epochs of fast locomotion in WT (t_(63)_=2.17, p=0.03) but not in β2^−/−^ mice (t_(50)_=-0.82, p=0.41, Fig. 7A), adding evidence to an involvement of β2*nAChRs in the dopamine encoding of exploration.

This prompted us to assess whether VTA pDAn activity already signaled novelty-induced exploration (Schiemann et al., 2012), in a new open-field without reward, i.e. before learning of the ICSS task (see Methods). Average firing frequency and %SWB did not differ in a novel and familiar environment (Supplementary Fig. 10), ruling out a pure dependence of exploratory locomotion on spontaneous DA activity. However, pDAn bursting activity during slow locomotion (Fig. 7B) was higher than during fast locomotion in WT (t_(51)_=-5.49, p<0.001), but not in β2^−/−^ mice (W_(30)_=119, p=0.15), indicating a similar encoding of locomotor exploration by the bursting activity of neurons in the contexts of novel environments and uncertain rewards (Fig. 7B and A).

## DISCUSSION

### Nicotinic acetylcholine receptors control dopamine firing on different timescales

nAChRs are known to affect the transition of dopamine firing from regular spiking to bursting (Dautan et al., 2016; Grace et al., 2007). Here we delved further on how this cholinergic modulation relates to behavior. Bursting is observed during phasic dopamine activity (Fiorillo et al., 2003; Jin and Costa, 2010; Lak et al., 2014; Redgrave and Gurney, 2006; Schultz, 2007), usually described as a reward prediction error (Glimcher, 2011; Schultz, 2007; Sutton and Barto, 1998), while tonic fluctuations have been related to most of the other functions ascribed to dopamine, e.g. exploration (Frank et al., 2009; Humphries et al., 2012), locomotion (Niv, 2007), and encoding of reward uncertainty (Fiorillo et al., 2003). However, the mapping between behaviors and dopamine dynamics remains unclear (Barter et al., 2015; Berridge, 2012; Howe and Dombeck, 2016; Lak et al., 2014), and it is thus crucial to distinguish the firing patterns (regular or bursting) from the timescale (average activity or event-related) considered. When considering average activity in anesthetized animals or at rest, bursting occurs spontaneously in dopamine cells (Faure et al., 2014; Mameli-Engvall et al., 2006; Schiemann et al., 2012; Schultz, 2007), but this bursting propensity is distinct from phasic, stimulus- or action-locked activity. Conversely, averaging stimulus-related activity from repeated trials might blur the analysis of firing patterns (Fiorillo et al., 2005; Niv et al., 2005).

Hence, we based our analysis on a definition of phasic activity that translates into large dopamine release (Faure et al., 2014; Grace et al., 2007), i.e. both bursting (i.e. short inter-spikes intervals) and synchronous among dopamine cells. We used this distinction to investigate the functional roles of dopamine neurons, and their upstream control by nicotinic acetylcholine receptors, on different timescales: on average (at rest or during the exploration of a novel environment), or at behaviorally-relevant timings during learning and decision-making tests.

First, we found that bursting onsets were spontaneously synchronized in awake mice at rest, suggesting that active mechanisms control the collective dynamics in the VTA (Eshel et al., 2016; Joshua et al., 2009). We suggest, among others, a nicotinic mechanism, as dopamine cells from β2^−/−^ mice did not display spontaneous synchrony between bursting onsets. However, contrary to previous observations in anesthetized animals (Mameli-Engvall et al., 2006; Maskos et al., 2005), we did find (non-synchronized) spontaneous bursting in dopamine cells from β2^−/−^ mice. This is consistent with bursting in dopamine cells arising from both cholinergic and glutamatergic inputs to the VTA (Faure et al., 2014). In awake animals, but not in anesthetized ones, excitation caused by glutamatergic inputs and muscarinic acetylcholine receptors would be sufficient to induce bursting activity in dopamine neurons. VTA β2*nAChRs would rather mediate a coordinated switch in dopamine cells towards a common excitable state (Faure et al., 2014; Grace et al., 2007), thus promoting synchronized bursting activity.

Second, we also show that the learning-induced shift of phasic dopamine activity from reward to reward-predictive stimulus (Fiorillo et al., 2003; Lak et al., 2014) or action (Jin and Costa, 2010; Wassum et al., 2012) relies on a re-organization of spiking towards synchronous bursting activity, with average firing frequency remaining stable. As reinforcement-learning theories would instead predict an addition of action potentials representing an increase in reward prediction, this suggests that both average VTA activity and its coordination among dopamine cells are tightly regulated. The same shift of phasic bursting occurs in dopamine cells from β2^−/−^ mice, but with less synchronous bursting and an increase in average firing frequency. This implicates β2*nAChRs not only in the spontaneous coordination of VTA activities, but also in the refinement of collective dynamics upon learning.

Third, our data indicate a role for β2*nAChRs in the encoding of uncertainty and exploration by both tonic and phasic dopamine firing. Before discussing (see below) how it furthers our understanding of uncertainty-seeking and exploration, these results show that nicotinic-cholinergic modulation of dopamine activity operates on different timescales, affecting slowly-fluctuating dopaminergic tone but also fast, coordinated firing at behaviorally-relevant timings. While acetylcholine has been implicated in the transition of dopamine firing from regular spiking to bursting pattern (Dautan et al., 2016; Grace et al., 2007), here we show that nicotinic receptors can affect such transition on a fast timescale (on the order of a hundred of milliseconds), and thus may impact immediate decision-making.

### What roles for the cholinergic control of dopamine in action, choices and exploration?

While there is a general agreement that dopamine is necessary for actions aimed at obtaining potential rewards (Berridge, 2012; McClure et al., 2003; Schultz, 2007), the contributions of dopamine to decision-making remain under investigation (Barter et al., 2015; Berridge, 2012; Howe and Dombeck, 2016; Nicola, 2010; Salamone and Correa, 2012; Schultz, 2007). In the reinforcement-learning framework, phasic dopamine signals a reward prediction error, which indirectly affects future choices by updating the cached values of options. However, there is an ever-growing debate about the direct role of dopamine at the time of choice (Glimcher, 2011; Schultz, 2007). This motivational and/or motor signal has been linked to phasic dopamine (Barter et al., 2015; Berridge, 2012; Howe and Dombeck, 2016) or to slower variations (Niv, 2007; Schultz, 2007). Even though we did not use “online” causal tools such as optogenetics (which can lead to erroneous inference about neural-behavior relations (Jazayeri and Afraz, 2017)), our correlational results indicate that phasic, rather than tonic, dopamine activity occurring prior to the initiation of goal-directed actions (Eshel et al., 2016; Jin and Costa, 2010; Joshua et al., 2009) signals both immediate choices (Morris et al., 2006) and kinetic variables of the goal-directed action (Barter et al., 2015; Howe and Dombeck, 2016). Some variants of reinforcement learning (“incentive salience” variant of actor-critic models (McClure et al., 2003)) offer an interpretation of the direct effects of phasic dopamine, by proposing that the reward prediction error affects the probability and latency to perform an action (McClure et al., 2003). However, we found that the phasic activity of VTA dopamine cells at the initiation of action sequence correlated with the kinematics of the whole sequence rather than just the latency to initiate an action. Our data thus suggest a more extended role of dopamine in motor control (Rigoux and Guigon, 2012).

Classic accounts also relate exploration, both in terms of decision-making and locomotion, to tonic, rather than phasic, dopamine (Frank et al., 2009; Humphries et al., 2012). Exploration has been linked to the excitability of dopamine cells (Naudé et al., 2016; Schiemann et al., 2012) or downstream signaling (Beeler et al., 2010; Frank et al., 2009; Humphries et al., 2012), but direct experimental observations of which firing patterns encode exploration were still lacking (Humphries et al., 2012; Naudé et al., 2016). We found that phasic activity correlated with uncertainty-seeking (discussed in-depth below) but that the modulation of tonic activity by uncertainty did not predict choices, and occurred at the end of the trial, presumably long after the decision process. Hence, it is doubtful that tonic encoding of uncertainty is directly implicated in choices under uncertainty, but may instead affect attentional processes (Fiorillo et al., 2003). We also observed that dopamine phasic activity at the beginning of trials could signal the distance travelled during long, exploratory trials. On a longer timescale, the extent of locomotor exploration was related to the modulation of bursting in dopamine cells when the animal was at low speed, rather than avergae bursting propensity. Hence, we overall show that locomotor exploration and uncertainty-seeking are both signaled by the phasic activity of dopamine cells. Our data add to the growing catalogue of functions (motor control (Barter et al., 2015; Howe and Dombeck, 2016), here exploration and uncertainty) long thought to be related to tonic dopamine (Niv, 2007; Schultz, 2007), that have proven signaled instead by fast, precisely timed, transitions towards phasic dopamine activity. In this framework, the disruption in the encoding of exploration by dopamine neurons from β2^−/−^ mice shows that cholinergic signaling can affect precisely-timed transitions of dopamine cells towards bursting, henceforth affecting decision-making on a short timescale.

### Cholinergic control of dopamine implements an uncertainty bonus

We also define how this fast effects of nicotinic receptors modulate dopamine encoding of decision variables. Together with previous studies, we show that phasic dopamine firing not only encode expected reward but also other reward attributes, such as risk (reward uncertainty (Fiorillo, 2011; Fiorillo et al., 2003; Lak et al., 2014; Stauffer et al., 2014)) or advanced information about upcoming rewards (Bromberg-Martin and Hikosaka, 2009), and non-reward attributes, such as novelty or sensory surprise (Redgrave and Gurney, 2006). Rather than a “reward-only” prediction error, which would encode the objective properties of a reward (e.g. probability, magnitude), dopamine neurons would instead signal the subjective value of a choice (Lak et al., 2014; Stauffer et al., 2014). The “subjective utility” theory remains however agnostic about the determinants of one’s preferences, i.e. it relates preferences for e.g. a type of food or for risky rather than safe option to dopamine firing, but does not explain what renders dopamine neurons activated by risk or uncertainty. Here we suggest how uncertainty can affect the subjective value of a choice, in terms of normative explanation and neural mechanisms.

We sought to explain the irrational preferences displayed by the animals in our setup: they display equal preference for the 50% and 100% options, even though the number and intensity of ICSS are the same for both outcomes. Using computational modeling (Daw et al., 2006; Funamizu et al., 2012; Naudé et al., 2016), we were able to extract the subjective value of uncertainty from animals’ choices, and to relate it with increases in phasic dopaminergic activity, on top of the encoding of expected reward. In the framework of phasic dopamine signaling a reward (or utility) prediction error, this means that the value of uncertainty is added to that of the expected reward, in the form of “bonus” value (Kakade and Dayan, 2002). Formalizing uncertainty-modulation of dopaminergic activity as a bonus explains suboptimal behavior as a motivation to obtain information about the current status of an unpredictable outcome (Anselme et al., 2013; Dayan, 2012; Vasconcelos et al., 2015), even if such information cannot be used to maximize reward in the future. Our task parameters, i.e., small stakes (there is no actual risk to lose, but just a risk not to win) and short delays between series of repeated gambles, are known to induce uncertainty-seeking (Heilbronner, 2013). In our set-up, the relative weight of uncertainty compared to expected reward was even close to one, meaning that mice priced (useless) information as much as rewards, reminiscent of playful or gambling behaviors (Oudeyer, 2007).

We also propose a neural mechanism for uncertainty-seeking. As VTA dopamine neurons from β2^−/−^ mice encode reward expectation and directness but not uncertainty or exploration, these different types of motivation arise from distinct dynamical processes at the level of the VTA (presumably also involving VTA GABA neurons (Tolu et al., 2012)), with exploration-related dopamine activity involving nAChRs. This implication of cholinergic mechanism in information seeking suggest that acetylcholine might signal uncertainty, as proposed by computational theories (Yu and Dayan, 2005), or that the coordinated switch in dopamine cells excitability gates (Grace et al., 2007) uncertainty signals generated elsewhere, e.g. in the cerebral cortex. This implication of β2-nAChRs in uncertainty-modulation of phasic activity (i.e. β2-nAChRs would be necessary for the integration of the uncertainty bonus) may explain the role of nicotinic receptors in probability discounting and henceforth in decisions under uncertainty (Fobbs and Mizumori, 2014). Overall, our results suggest that pontine cholinergic modulation, far from being just a slowly fluctuating tone, can affect dopamine firing on a fast timescale, thereby instantly biasing animals towards information-gathering.

## Author contribution

J.N. and P.F. designed the study. S.D. and S.T. designed the microdrives for electrophysiology and performed open-field “novelty” experiments. J.N. and C.P-S performed the “ICSS multi-armed bandit” experiments. J.N. S.D. S.T. and P.F. analyzed the data. U.M. provided genetically-deleted mice. J.N. and P.F. wrote the manuscript with inputs from S.D. and S.T.

## Acknowledgments

We thank Alexandre Mourot, Emmanuel Guigon and Camilla Bellone for comments on the manuscript. This work was supported by the Centre National de la Recherche Scientifique CNRS UMR 8246, the Fondation pour la Recherche Médicale (FRM, Equipe FRM DEQ2013326488 to P.F., fellowship FDT20150532705 to S.D.), the Agence Nationale pour la Recherche (ANR Programme Blanc 2012 for P.F.), the Fédération pour la Recherche sur le Cerveau (FRC 2012 to P.F.), the Bettencourt Schueller Foundation (Coup d’Elan 2012 to P.F.), the French National Cancer Institute Grant TABAC-16–022 (to P.F. and U.M.), Pierre et Marie Curie University (Programme Emergence 2012 to J.N. and P.F.; PhD fellowship to S.D.), Pierre et Marie Curie University (to S.T. and S.D.). P.F. laboratory is part of École des Neurosciences de Paris Ile-de-France RTRA network. P.F. is a member of LabEx Bio-Psy and of DHU Pepsy.

## METHODS

### 1. Animals

Experiments were performed on 14 wild-type (WT; C57BL/6J) and 11 knockout SOPF HO ACNΒ2 (β2^−/−^) male mice obtained from Charles Rivers Laboratories Frances. β2^−/−^ mice were generated as described previously (Picciotto et al., 1995). Even though WT and β2^−/−^ mice are not littermates, the mutant line was generated more than 20 years ago, and has been backcrossed more than 20 generations with the wild-type C57BL6/J line, and is more than 99.99% C57BL/6J. All experiments were performed on mice between 2 and 4 months of age. Mice were housed in cages containing a maximum of 4 animals, in a 12 h light/dark cycle and temperature-controlled room (22 +/− 1 °C) with food and water available ad libitum. Sample sizes in this study are similar to those generally employed in the field and were not pre-determined by a sample size calculation. 4 wild-type and 4 β2^−/−^ mice were excluded from the analysis due to improper electrode implantation (no dopamine neurons were found in these mice). All experiments were undertaken in compliance with French laws on animal experimentation, the directives of the European Community 219/1990 and 220/1990. The protocol was approved by the Committee on the Ethics of Animal Experiments of the University of Pierre et Marie Curie (Permit Number: 01438.01).

### 2. Drive and electrodes

Hand-made poly-electrodes (bundle of 8 electrodes, “octrodes”) were obtained by twisting eight polyimide-insulated 17 μm Nickel-Chrome wires (A-M SYSTEMS, USA). The use of eight channels relatively close together allows for a better discrimination of the different neurons. Before implantation and recording, the octrodes were cut at suitable length and gold-plated (gold plating solution, Neuralynx; Bozeman, USA) to lower their impedance to 500–800 KOhms and improve the signal-to-noise ratio. The free ends of the octrodes are connected to the holes of EIB-18 (electrode interface board, Neuralynx) and fixed with pins. Two octrodes were placed in a cannula (90-micron diameter) to guide the octrode to the brain and the upper part of the octrode without damaging its tip. We manufactured a microdrive system (home-made 3D conception and printing) consisting of a main body, on which is mounted the EIB, and a driving screw, with a sliding part comprising the octrode-containing cannula. This microdrive allowed moving through the VTA in order to sample neuronal populations Finally, a bipolar stimulation electrode for the IntraCranial Self-Stimulation (ICSS) was also fixed to the EIB (in the stimulation ports).

### 3. Surgery

After induction of anesthesia with a gas mixture of oxygen (1l/min) and 3% isoflurane (Vetflurane^®^, Virbac), the mouse was placed in a stereotaxic frame (David Kopf). The cranial bone was exposed by a midline incision of the skin. The skull was then drilled and recordings electrodes were placed just above the VTA (coordinates: 3.2±0.1 mm posterior to the Bregma, 0.5±0.1 mm lateral and 4.0±0.1 mm deep from the surface of the brain (Paxinos and Franklin, 2004)). From this starting position, electrodes were lowered (75μm steps) during the experiment until a depth of 5.0 mm was reached. Monopolar ground electrodes were laid over the cortical layer of the cerebellum and the olfactory bulb. Stimulation electrode for ICSS was implanted in the medial forebrain bundle (MFB, 1.4 mm posterior to the Bregma, mediolateral= ±1.2 mm lateral and 4.8 mm deep from the surface of the brain). Dental acrylic cement was used to fix the main body of the microdrive to the skull during the surgery. After surgery, an antiseptic (Povidone37 iodine solution) and a local anesthetic (lidocaine ointment) were applied in areas where the scalp was incised. Animals recovered until regaining pre-surgery body weight, and at least two weeks.

### 4. Neuronal recordings and characterization of dopamine neurons

Recordings of extracellular potentials were performed using a digital acquisition system (Digital Lynx SX; Neuralynx) together with the Cheetah software. Broadband signals from each wire were filtered between 0.1 and 9000 Hz and recorded continuously at 32 kHz. To extract spike timing, signals were band-pass-filtered between 600 and 6000 Hz and sorted offline. Spike clustering was cross-validated by using both SpikeSort3D (Neuralynx) and custom-written Matlab (The Mathworks) routines (spike-sorting codes are available upon request). The electrophysiological characteristics of VTA neurons were analyzed in the active cells encountered by systematically moving down the octrode-containing cannula. Extracellular identification of putative DA neurons was based on their location as well as on a set of unique electrophysiological properties that characterize these cells in vivo: 1) a typical triphasic action potential with a marked negative deflection; 2) a characteristic long duration (>2.0 ms) action potential; 3) an action potential width from start to negative trough >1.1 ms; 4) a slow firing rate (<10 Hz) with an irregular single spiking pattern and occasional short, slow bursting activity. Putative GABA neurons were characterized by a characteristic short duration of action potential from start to negative trough (<1.0 ms), and a high firing rate (>12Hz). D2 receptors (D2R) pharmacology was also used for confirmation of DA neuron identification: after a baseline (10 min) and a saline (dose, 10 min) injection, quinpirole (1mg/kg,, D2R antagonist) was injected (30 min recording), followed by an eticlopride (D2R agonist) injection (1mg/kg, 30 min recording). Since most DA, but not GABA neurons, express inhibitory D2 auto-receptors, neurons were considered as pDA neurons if quinpirole induced at least 30% decrease in their firing rate, while eticlopride restored firing above the baseline. Nevertheless, as continuous D2 pharmacology could have affected both baseline DA neurons firing and decision-making (Fobbs and Mizumori, 2014), we allowed the mice to recover two days after this experiment. We thus performed pharmacological confirmation (1) when first encountering a putative DA neuron in a given mouse or (2) at the end of the week if at least one putative neuron was present during the behavioral experiment. Neurons were considered DA only if they responded to the pharmacology, or if they presented electrophysiological characteristics defined above and were recorded between two positive pharmacological experiments.

### 5. Immunohistochemistry

Immunolabeling allowed to assess the TH-positive phenotype of the neurons surrounding the electrode trace and was performed as follows. All neurons from mice in which the electrode trace was detected outside the VTA were excluded from the analysis. Following the death of the mice, brains were rapidly removed and fixed in 4% paraformaldehyde. After at least 3 days of fixation at 4 °C, serial 60-μm sections were cut from the midbrain with a vibratome. Free-floating VTA brain sections were incubated 1h at 4°C in a blocking solution of phosphate-buffered saline (PBS) containing 3% Bovine Serum Albumin (BSA, Sigma; A4503) and 0.2% Triton X-100 and then incubated overnight at 4°C with a mouse anti-tyrosine hydroxylase antibody (TH, Sigma, T1299) at 1:200 dilution in PBS containing 1.5% BSA and 0.2% Triton X-100. The following day, sections were rinsed with PBS and then incubated 3 h at 22–25°C with Cy3-conjugated anti-mouse secondary antibodies (Jackson ImmunoResearch, 715–165–150) at 1:200 dilution in a solution of 1.5% BSA in PBS. After three rinses in PBS, slices were wet-mounted using Prolong Gold Antifade Reagent (Invitrogen, P36930). Microscopy was carried out with a fluorescent microscope, and images captured using a camera and ImageJ imaging software.

### 6. Behavioral data acquisition

Locomotor activity and decision-making (ICSS choice task, see below) were both recorded in a 0.8-m diameter circular open-field. Experiments were performed using a video camera, connected to a video-track system, out of sight of the experimenter. A home-made software (Labview National instrument) tracked the animal, recorded its trajectory (20 frames per s) for 5 min and sent TTL pulses to the ICSS stimulator when appropriate (see below). Spontaneous activity (behavioral and neuronal) was obtained in the mouse home-cage (“familiar environment”) for 10 minutes prior to starting the open-field experiment. The “new environment” corresponds to the open-field, the first time the animals were exploring it, for 30 minutes. Animal trajectories were smoothed using a triangular filter. Time-to-goal measures the duration of one trial between one ICSS location and the next one. The speed profile corresponds to the instantaneous speed as a function of time. Maximal speed and time-to-maximal speed were detected for each trial after removing the first second after the last ICSS location, which corresponds to the duration where early bursting firing activity was computed (see below). In the new environment and in the long, exploratory trials of the probabilistic setting (Fig. 6,7), behavior was decomposed according to animal’s speed between navigation and exploration epochs using a dual threshold criterion (Maskos et al., 2005; Maubourguet et al., 2008). The behavior was considered as to change from “exploration” to “navigation” if and only if speed crossed the high threshold (12 cm.s-1) while the behavior was considered to change from navigation to exploration only if its speed crossed the low threshold (speed < 6 cm.s-1), avoiding spurious navigation and exploration phases of small duration.

### 7. Markovian decision problem by ICSS conditioning

Details for this experiment can be found in a previous publication (Naudé et al., 2016). Briefly, three explicit square locations, marked on the floor, were placed in the open field, forming an equilateral triangle (side = 50 cm). Each time a mouse was detected (by its centroid) in the area of one of the rewarding locations (area radius = 3 cm), a 200-ms train of twenty 0.5-ms biphasic square waves pulsed at 100 Hz was delivered. Experiments were performed using a video camera, connected to a video-track system, out of sight of the experimenter. A home-made software (Labview National instrument) tracked the animal, recorded its trajectory (20 frames per s) for 5 min and sent TTL pulses to the ICSS stimulator when appropriate. Animals received stimulations only when they alternate between rewarding locations. The training consisted of a block (10 daily sessions of 5 min) in a certain setting where all locations were associated with an ICSS delivery. ICSS intensity was adjusted so that mice self-stimulated between 50 and 150 times per session at the end of the training (ninth and tenth session). Current intensity was subsequently maintained the same throughout the probabilistic setting. The test phase consisted of a block (10 daily sessions of 5 min) assessing choice organization under a probabilistic setting, by associating each location with a different probability of ICSS (25%, 50% and 100%, randomly assigned for each mouse). Reward probabilities were pseudo-randomly assigned to each location for each mouse. Animals underwent an additional 5-sessions blocks of probabilistic settings with different probability sets (25%, 50% and 75%; 25%, 75%, and 100%; 50%, 75% and 100%), pseudo-randomly assigned, if dopamine neurons were still recorded in these animals. From 14 WT and 10 β2^−/−^ mice at the beginning of the study (i.e. training), 10 WT and 7 β2^−/−^ mice had dopamine neurons in the probabilistic sessions.

### 8. Analysis of decision-making

Animal trajectories were smoothed using a triangular filter. Time-to-goal measures the duration of one trial between one ICSS location and the next one. The speed profile corresponds to the instantaneous speed as a function of time.

Analysis of decision-making was based on a previous study (Naudé et al., 2016). In short, we expressed behavioral data as a series of choices between rewarding locations (labeled A, B, C). We only considered choices made in an interval of 10s after visiting the previous rewarding location. This experiment implements a Markovian Decision Process (MDP (Sutton and Barto, 1998)) consisting of three states (A, B, C), corresponding to each rewarding locations, and a transition function, corresponding to the proportions of choices in the three gambles. The repartition is defined as the proportion of states visited by the animal during a session. The transition matrix describes the proportion of transitions from one state to another. Because animals receive stimulations only when they alternate between rewarding locations, there is no repetition of states in the sequence and the 3 × 3 transition matrix has null diagonal elements.

We modeled mice decision strategies with a previously-published computational model, an uncertainty-sensitive softmax rule, henceforth called the “uncertainty model”. Because mice could not return to the same rewarding location, they had to choose between the two remaining states (rewarded locations). The uncertainty model determined the probability P_i_ of choosing the next state i, as a function of a decision variable. The softmax choice rule is:

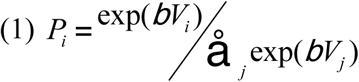
where β is an inverse temperature parameter (“decision noise”) reflecting the sensitivity of choices to the difference between decision variables. In our model embedding an exploration bonus, the decision variable (the value of an option) depends on both expected reward and uncertainty (Daw et al., 2006; Frank et al., 2009; Funamizu et al., 2012).

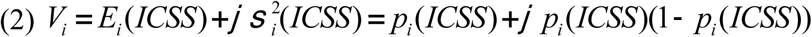

This compound value is then nested in the softmax choice rule. We fitted the free parameters (β, φ) maximizing the respective likelihood of the observed choices c at all trials t for each mouse separately, using the population fit (fit of the average choice probabilities) as initial conditions.

### 9. Electrophysiology analysis

Both spontaneous activity and behavior-related neuronal activity were analyzed offline with Matlab custom-written codes. Spontaneous DA cell firing was analyzed with respect to the average firing rate and the percentage of spikes within bursts (%SWB, number of spikes within bursts, divided by total number of spikes). Bursts were identified as discrete events consisting of a sequence of spikes such that: their onset is defined by two consecutive spikes within an interval <80 ms and they terminated with an interval >160 ms. Phasic activity is defined as spikes falling into bursts, while tonic activity comprises spikes outside bursts. Synchrony between pairs of neurons was computed according to Kreuz et al.(Kreuz et al., 2013) definition, with SPIKY matlab code. Briefly, spike synchronization measures the fraction of coincidence between two spike trains. This measure is comprised between 0 (no coincidence) when the spikes trains are most dissimilar and 1 (each spike has one matching spike in the other train) when they are similar. This method is parameter- and scale-free, because the coincidence window is adapted to the local spike rate. It is also time-resolved, which allows assessing synchrony during trials. Peri-event time histograms were constructed based on 1 ms-bins rasters, convolved with a Gaussian kernel (width = 50ms) corresponding to the same event. This event corresponded to the reward in the certain setting. In the probabilistic setting, different peri-event time histograms were computed, corresponding to each reward probability of the next location visited by the animal (e.g. 25%, 50%, 100%), centered on animal start (i.e. the previous location occupied by the mouse) or arrival at the next location. As DA neurons may fire action potentials at frequencies ranging from 1 to 10Hz, to avoid bias towards responses of high firing neurons, firing frequencies from the peri-event time histograms were normalized for each neuron by its average firing frequency, and then averaged over the population. Neurons were considered as having an early bursting activity at the beginning of the between-location movement if their firing frequency was 2 times above baseline for 20 consecutive time bins (Jin and Costa, 2010) in the first quarter of the movement. For between-location plots (labeled “linearized time between locations” in Fig. 2), DA firing activity was binned (1000 bins) from 0 to 100% of the duration from the previous location to the target location, and then normalized by average frequency and averaged over the population. Average firing as a function of linearized time was only used for display purposes and was not analyzed. Early bursting frequency and ramping frequency were computed on direct trials (>5s). Early bursting frequency was computed on the first 500ms following the last location, whether the animal received an ICSS reward (hence, after the 200 ms duration of the ICSS) or not (in this case, 200 ms corresponding to reward omission were also skipped). Ramping frequency was computed on the last second preceding the target location.

### 10. Statistical analysis

No statistical methods were used to predetermine sample sizes, which are comparable to many studies using similar techniques and animal models. We used a pseudo-randomization procedure, in the sense that in the behavioral experiments, precise parameters (for example, reward probabilities) were pseudo-randomly assigned to each rewarding location for each mouse. Behavioral and electrophysiological data were analyzed and fitted using Matlab (The MathWorks). Data are plotted as mean ± s.e.m. Total number (n) of observations in each group and statistics used are indicated in figure captions. Classically comparisons between means were performed using parametric tests (Student for two groups, or ANOVA for comparing more than two groups) when parameters followed a normal distribution (Shapiro test P > 0.05), and non-parametric tests (here, Wilcoxon or Mann-Whitney) when this was not the case. Homogeneity of variances was tested preliminarily and the t tests were Welch-corrected if needed. Multiple comparisons were Bonferroni corrected. All statistical tests were two-sided. P > 0.05 was considered to be not statistically significant.

